# Genetic variation in a plant-parasitic nematode is associated with gene expression changes conserved across evolutionarily distant host plants

**DOI:** 10.64898/2026.01.07.698247

**Authors:** Dahlia M. Nielsen

## Abstract

A hallmark of the interactions between endoparasites (parasites that live within plant tissue) and their hosts, is the parasites’ ability to hijack host gene expression to support their own existence. While a number of studies have been performed to identify plant genes that are recruited in this process, there is much that remains unknown. One mechanism by which these connections can be explored is using a traditional genetics approach – identifying connections between genetic variation and phenotype variation. To that end, we leverage experiments examining tissue from three host plants, each infected with one of two genetically distinct strains of the endoparasite *Meloidogyne hapla*. Using gene expression profiles, we identify plant genes that differ based on the genetic background of the parasite, and show that these signals are common across plant hosts. These results indicate that the plant genes recruited by the parasite differ according to the parasite’s genetic background, and that these alternative patterns are conserved across evolutionarily distant hosts. We also leverage public data to identify conserved host expression differences when comparing *M. hapla*-infected roots to uninfected root tissue. Finally, we demonstrate that parasite gene expression varies based on the parasites’ genetic background, and these signals are also conserved when inhabiting different host species.

## INTRODUCTION

One of the astonishing evolutionary adaptations endoparasites have in common is their ability to hijack host gene expression, cellular processes, and overall development to create an environment beneficial for feeding and evading host defenses (Favery et al. 2020). Various studies have been performed to identify plant genes involved in these interactions (Liu et al. 2007; Simonetti et al. 2009; Kawazu et al. 2012; Aditya et al. 2015; Dowd et al. 2017; Bournaud et al. 2018), including comparisons across different species of hosts and parasites (Takeda et al. 2019). However, much yet remains unknown about the plant genes involved, the mechanisms by which they are recruited, or how development is altered based on the hijacked pathways.

Root knot nematodes (RKN; *Meloidogyne* spp.), a ubiquitous parasite of vascular plants, are one such endoparasite. These have been a focus of study due to their substantial worldwide negative agricultural impact (Jones et al. 2013). Like other endoparasites, RKN coerce hosts into establishing and maintaining permanent feeding sites (Gheysen and Mitchum 2011; Favery et al. 2016). Central to these sites is the formation of host cells termed Giant Cells (GC) that serve as the nutrient source for the nematode (Bird and Loveys 1975). While successful development of GC is common to *Meloidogyne* spp., substantial variation has been observed in field isolates, including in host responses elicited by the nematodes (Janssen et al. 1997; Trudgill and Blok 2001; Melakeberhan and Wang 2012). One expected source of these phenotypic differences is population- and species-level genetic variation. As it is reasonable to assume that genetically variable RKN lines take different paths to establish an infection, connecting genetic variation in the parasite to specific host responses has the potential to elucidate host genes and host pathways that are usurped by the nematode for its own gain.

We have previously shown that by inoculating isogenic Medicago (*Medicago truncatula*) plants with *Meloidogyne hapla* lines from a mapping population (derived from a bi-parental cross), we could tie expression networks of plant and parasite genes to specific loci in the nematode genome (Guo et al. 2017). Results from this cross-species genetic mapping study using Medicago were mirrored in RKN-infected tomato plants.

Here we perform a broader investigation of host responses. We combine three experiments performed previously to examine how genetic variation in the RKN *M. hapla* affects plant host response, each using a different host species. For each of these experiments, we leveraged two genetically divergent isogenic strains of *M. hapla* (VW9 and LM). The three experiments used different host plants: Medicago (*Medicago truncatula*), soybean (*Glycines max*), and tomato (*Solanum lycopersicum*). Each experiment followed the same experimental design: for each host species, isogenic plants were used; half of the plants were infected with VW9, and half with LM. Other environmental conditions were maintained to be consistent across experiments. For each experiment, differential expression was tested between VW9-infected and LM-infected plant tissue (galls). If a gene was identified as being differentially expressed, the direction of the effect was also recorded (whether expression was higher in VW9-infected plants than LM-infected plants, or vice versa). Because infected tissue was assayed, expression patterns could be assayed for both the plant host and the nematode parasite.

These experiments allowed us to examine the effect of genetic variability of the parasite on gene expression levels of the host. Our premise was that genes predicted to be orthologous across species would share similar expression responses. Significant overlap of differentially expressed genes across host plant species provides an indication that expression differences between VW9- vs LM-infected plants are conserved across evolutionary distance. We could additionally test nematode gene expression for differences between strains, and determine whether patterns in pathogen gene expression varied based on the species of their host. Finally, with some caveats, we were able to compare host plant expression between infected and uninfected tissue downloaded from public databases (Huang and Schiefelbein 2015), and to assess the conservation of these signals across host species.

## RESULTS

Data from three experiments were used. In each experiment, replicate plants from a single accession were assayed, half of which were infected with *M. hapla* strain VW9, and half infected with strain LM. Each experiment followed the same infection and sample collection protocol (see Methods for details). Three weeks post-infection, infection sites (root galls) were dissected and tissue processed for RNA-Seq. By using infected tissue, RNA from both the host and the parasite was jointly collected. Sequence reads generated were aligned to a concatenated reference genome containing both the *M. hapla* and the respective plant genome sequences. Once sequence reads were aligned, their species of origin (plant or nematode) could be determined based on the alignment results. Normalization and analysis of RNA-Seq data was performed separately for plant reads and nematode reads, as were tests for differential expression (see Methods for details on data processing and analysis). In all experiments, comparisons of expression were made between VW9-infected and LM-infected plant tissue. If a gene was identified as being differentially expressed, the direction of the effect was also recorded (whether expression was higher in VW9-infected plants than LM-infected plants, or vice versa).

### Genetically distinct lines of *M. hapla* elicit broadly different transcription responses in their plant hosts

We first examined each host separately, comparing gene expression of plants infected with the VW9 line of *M. hapla* to plants infected with the LM line. In the Medicago experiment, 950 genes were identified as being differentially expressed. In soybean, 2,341 genes were identified, and in the tomato experiment, 1,847 genes were identified. For all host species, the results with the most statically significant p-values were MADS-box transcription factor (TF) genes.

### Gene expression responses to parasitic genetic variants are conserved across plant host species

Once genes that were differentially expressed between VW9- and LM-infected plants were identified for each host species, we were interested in comparing results across species. Our premise was that genes predicted to be orthologous across species would share similar expression responses. To test this, orthologous gene sets (orthogroups) were first generated using OrthoFinder (Emms and Kelly 2015, 2019). We then merged the orthogroup information with differential gene expression results for each host plant experiment.

We focused on orthologous sets containing expressed genes from more than one host species. For these, we examined whether significant differential expression results tended to cluster within orthogroups (in other words, if a gene is differentially expressed for one host species, is it more likely that orthologous genes from another host species are also differentially expressed). If this is the case, this is evidence that differences in host responses caused by genetic variation in the parasite is conserved across host species. To assess this, we performed both Fisher’s Exact tests (FET) for each pair of species. We found that differential expression tended to be conserved (FET p-values: Medicago-soybean, p=1.67e-29; Medicago-tomato, p=3.86e-17; soybean-tomato, p=5.23e-28).

Interestingly, while genes within orthologous gene sets tend to agree in terms of being differentially expressed, the direction of the expression change was surprisingly inconsistent. We identified 707 orthologous groups that contained differentially expressed genes from at least two host plant species. Of these 707 orthogroups, 423 (∼60%) displayed signals for which the direction of the effect was the same for all genes Significant results from the remaining 284 groups (∼40%) displayed signals that were reversed for some genes relative to other genes in the group.

Results for genes from the same species within orthogroups, however, did show strong consistency. There were 662 orthogroups that contained two or more differentially expressed genes from individual host species. For each host species, we selected two genes from each orthogroup (these were selected at random if more than two genes were differentially expressed for that species in that orthogroup). Using these gene pairs, we applied a FET for each host species to test for consistency of the direction of expression. All three host species showed significant agreement in the direction of differential expression between the gene pairs (Medicago, p=7.8e-12; soybean, p=1.8e-49, tomato, p=7.5e-12).

### Gene expression responses between infected and uninfected plants are conserved across host plant species

We were also interested in using our data to examine differential expression between infected and uninfected plants. To address this, we utilized data collected by Huang and Schiefelbein, 2015 (GEO accession number GSE64665). These investigators examined expression patterns of root development in seven plant species. For each species, they collected RNA-Seq data for three samples from each of three root zones: the root meristematic zone, the root elongation zone, and the root differentiation zone. Two of the species they used were the same species and accessions that we assayed: soybean (*G. max*, accession Williams 82) and tomato (*S. lycopersicum*, accession Heinz 1706). We downloaded their raw sequence reads for tomato and soybean to compare with our LM- and VW9-infected tissue.

There are various technical factors that can lead to genes displaying differential patterns of expression when examining data across studies. We were interested in determining whether we could identify biologically relevant signals in our comparisons. Sequence data was processed using the same pipeline as the infected tissue from our lab, and tests of differential expression were performed for all genes displaying detectable expression levels (see Methods for details). Results were then integrated based on orthogroup membership. All orthogroups that contained tomato and soybean genes that were expressed in either infected or uninfected samples were identified, and the results for the tests of differential expression for those genes were examined for agreement. Our rationale was that if the effects we identified were due to technical factors (caused by samples being collected and assayed in different labs), these effects would not be conserved between tomato and soybean data. Consistent results across species would indicate that the results are enriched for biologically relevant differences between infected and uninfected tissue. First, to explore the joined dataset of infected and uninfected plants, we selected pairs of expressed genes (one tomato and one soybean) from each orthogroup that contained such pairs (see Materials and Methods). Using orthogroups with these pairs, we performed a principal components analysis (PCA) to examine how samples clustered. Results are depicted in Figure 1. Based on the first two principal components (PC), samples are first clustered by species (PC1), then primarily by dataset type (infected vs uninfected; PC2).

**Figure 1.**
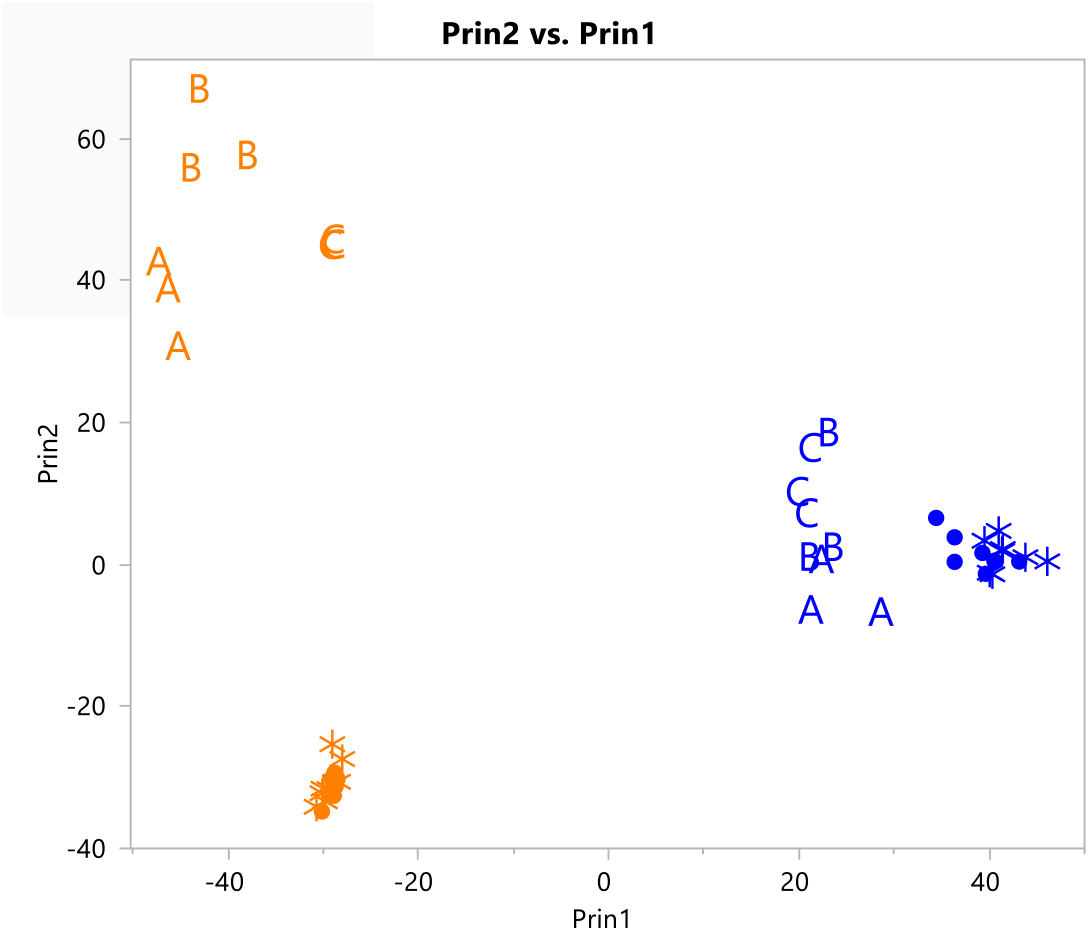
PCA results plot for merged *M. hapla*-infected and uninfected samples. Results are plotted for PC1 vs PC2. Tomato samples are in orange and soybean samples are in blue. The different uninfected root tissue types are designated by letters (A=root differentiation zone; B=root elongation zone; C=root meristematic zone). VW9-infected samples are demarcated with a circle, and LM-infected samples with an asterisk.

Differential gene expression results for orthogroups with one or more expressed gene for each of soybean and tomato were then compared using FET. For the 3,921 ortho groups containing both expressed tomato and soybean genes, agreement between the species was highly statistically significant (p=1.9e-55). This indicates that if one or more genes from one species within an orthogroup are differentially expressed, it is highly likely genes from the other species will also be differentially expressed.

### Differences in nematode gene expression between LM and VW9 are consistent across host plant species

We were interested in examining whether nematode genes were differently expressed depending on their own genetic background (VW9 vs LM strains of nematodes), and if so, if these differences were consistent when the nematodes inhabited alternate plant host species. Since RNA was extracted from infected root tissue, both plant and nematode gene expression can be assayed using the same samples. *M. hapla* gene expression was measured for LM and VW9 strains in all three plant hosts, and nematode genes were tested for differential expression. In this case orthogroup information is unnecessary, as the same parasite genes are measured for each plant background.

There were 8,888 *M. hapla* genes whose expression could be assayed and tested across the full experiment. Nematode expression was highly correlated between infections, with correlation coefficients of 0.93 (Medicago-soybean), 0.83 (Medicago-tomato) and 0.90 (soybean-tomato). Additionally, if a RKN gene was differentially expressed between LM nematodes and VW9 nematodes in one host species, it was highly likely to be differentially expressed in at least one other host (p < 1x10^-10^). In total, we find 93 RKN genes that are differentially expressed in all three plant host infections and 329 genes differentially expressed in two of the three host plant infections. In the vast majority of cases, the direction of the effect was consistent (if expression was higher in LM than VW9 in one plant host species, it was generally higher in the other plant host species). For the 93 genes that were differentially expressed between all three LM- and VW9-infected plant host species, 92 displayed a consistent effect (Figure 2).

**Figure 2.**
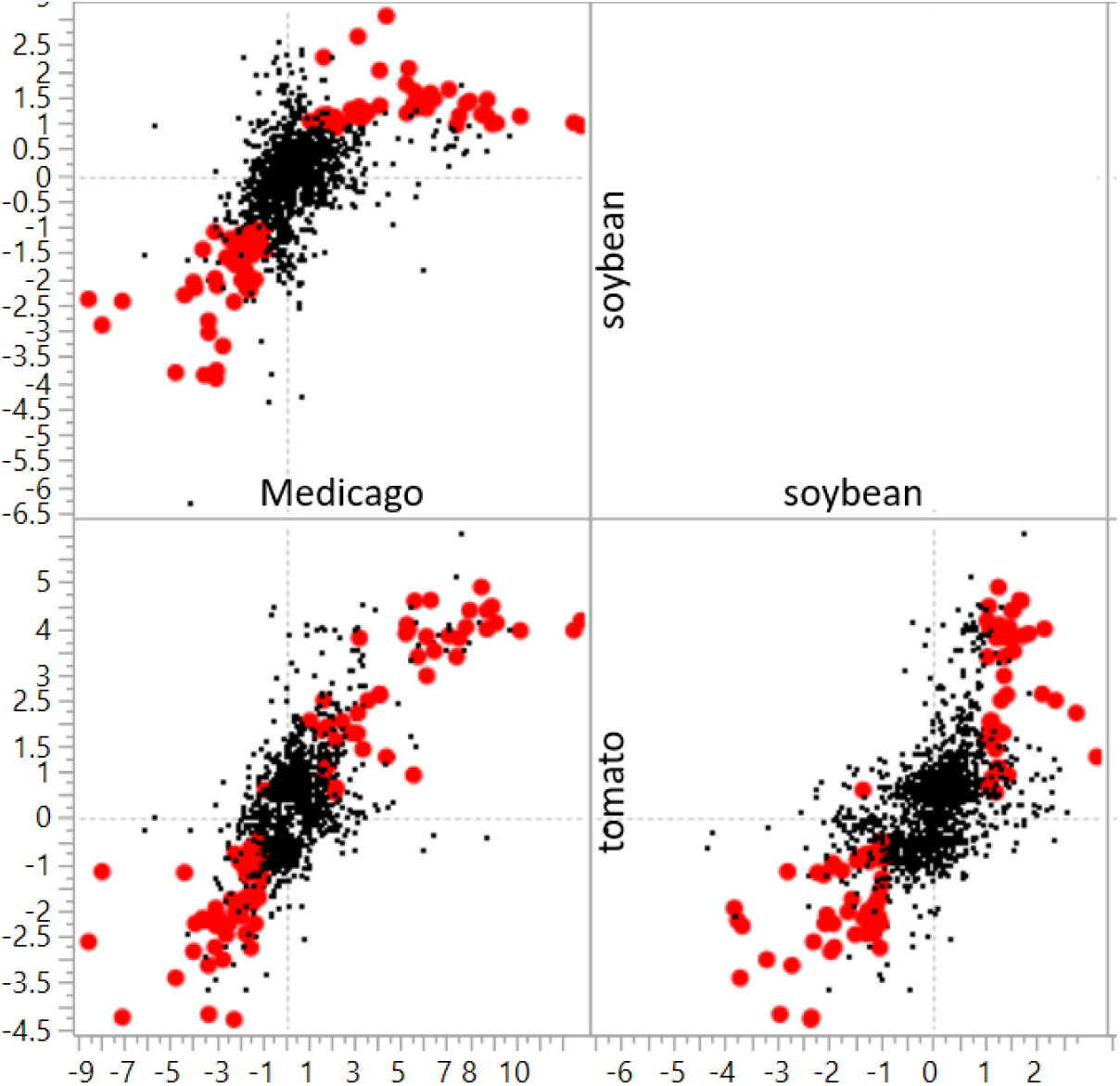
Correlated effects for RKN genes differentially expressed between strain LM and VW9 in each of the three host plant species. Each point is log2(fold change) for one RKN gene (LM-VW9). Red dots indicate genes that are differentially expressed in all three host plants. With the exception of one gene, the direction of the effect is the same for each gene (the directions match for Medicago and soybean, but are reversed for tomato).

## MATERIAL AND METHODS

### Experimental Procedures and Experimental Design

Experimental details related to the material used in these experiments, including nematode strains, plant species, and plant cultivars has been previously described (Guo et al. 2017). Each host plant experiment used for this study was originally performed as a stand-alone project. For each, isogenic plants were germinated and grown for two weeks before inoculation with either *M. hapla* strain VW9 or with strain LM. At three weeks post inoculation, resected sections of plant root (galls or root knots) harboring feeding nematodes were collected, and RNA (containing a mixture of plant and *M. hapla* transcripts) was extracted for RNA-Seq. Sequencing reads were generated and aligned to the concatenated reference of the *M. hapla* and appropriate plant genomes, enabling us to measure transcript abundance for both the RKN and its plant host. Sample sizes for each experiment were as follows. Medicago: four Medicago (Jemalong A17) plants were inoculated with *M. hapla* strain LM, and four were inoculated with hapla strain VW9. Tomato: eight tomato (Heinz 1706) plants were inoculated with LM strain *M. hapla* and eight were infected with VW9. Soybean: eight LM-infected soybean plants (Williams-82) were compared with seven VW9-infected plants (one of the VW9-infected plants did not provide sufficient quality RNA).

### RNA-Seq Data Processing and Analysis

RNA-Sequencing reads were aligned to the appropriate genome using HISAT2 version 2-2.2.1 (Kim, Langmead, and Salzberg 2015; Pertea et al. 2016; Kim et al. 2019) with default settings. Gene expression counts were generated from aligned sequence reads using HTSeq version 2.0.2 (Anders, Pyl, and Huber 2015; Putri et al. 2022). Counts were normalized and tests of differential expression were performed using edgeR (Robinson, McCarthy, and Smyth 2010; Y. Chen et al. 2025). Data were first read into an R data frame, and two groups constructed (one for each of VW9 and LM samples). Commands were then enacted as follows: dge = DGEList(counts = as.matrix(data), group = groups, genes = row.names(data)); dge = calcNormFactors(dge); dge = estimateCommonDisp(dge); de.com = exactTest(dge). Significance was determined (for each experiment separately) using the false discovery rate procedure of Benjamini and Hochberg (Benjamini and Hochberg 1995).

A gene was considered to be expressed if at least one group (VW9-infected plants, LM-infected plants, uninfected plants) in the comparison of interest had a median raw count of 10 sequence reads.

### Orthogroup Construction

To create the orthogroups, we used OrthoFinder version 2.5.4 (Emms and Kelly 2015). To aid in inference accuracy, we included predicted protein sequences from ten plant species, including the three examined here, representing plants from a broad evolutionary history. These included *Arabidopsis thaliana, Brassica rapa, Citrus sinensis* (orange), *Cucumis sativus* (cucumber), *Lotus japonicus, Mimulus guttatus* (monkeyflower), and *Vitis vinifera* (grape). Genome versions for each are given in Table 1.

**Table 1:** Genome versions of plant species used to identify orthogroups.

| Species | Genome resource |
| --- | --- |
| *Glycines max* | Glycine max Wm82.a4.v1 (Valliyodan et al. 2019; Schmutz et al. 2010; Goodstein et al. 2012) |
| *Medicago truncatula* | Medicago truncatula Mt4.0v1 (Tang et al. 2014; Young et al. 2011; Goodstein et al. 2012) |
| *Solanum lycopersicum* | Solanum lycopersicum ITAG4.0 (Hosmani et al. 2019; Goodstein et al. 2012) |
| *Arabidopsis thaliana* | Arabidopsis thaliana Araport11 (Cheng et al. 2017; Goodstein et al. 2012) |
| *Brassica rapa* | Brassica_rapa/V3.0 (H. Chen et al. 2022) |
| *Citrus sinensis* | C.sinensis_Hzau_v2.0 (citrusgenomedb.org) |
| *Cucumis sativus* | Chinese_long/v3/ (Q. Li et al. 2019) (cucurbitgenomics.org) |
| *Lotus japonicus* | Lotus japonicus Lj1.0v1 (H. Li et al. 2020; Goodstein et al. 2012) |
| *Mimulus guttatus* | Mimulus guttatus v2.0 (Hellsten et al. 2013; Goodstein et al. 2012) |
| *Vitis vinifera* | Vitis vinifera v2.1 (Jaillon et al. 2007; Goodstein et al. 2012) |

### Principal Components Analysis

Orthogroups containing expressed genes from both soybean and tomato were selected. If an orthogroup contained more than one gene for a given host species, one gene was selected at random. Pseudocounts (normalized gene expression counts) produced by edgeR (Robinson, McCarthy, and Smyth 2010; Y. Chen et al. 2025) were used to calculate the PCs. Analysis was performed using JMP (JMP® 2025).

### Analysis of Combined Results

All analyses involving the merged processed counts data and differential expression results were performed using JMP (JMP® 2025). This included Fisher’s Exact tests, Principal Components Analysis, correlation analyses, and all data manipulation such as the merging and filtering of datasets.

For analyses in which pairs of genes were selected from orthogroups (such as the PCA, where one expressed gene from each of soybean and tomato were selected), orthogroups were first identified that contained at least one such pair. If that orthogroup contained more than one gene to choose from to create the pair, genes were selected at random.

## DISCUSSION

Here, we combined results across three experiments designed to identify host plant genes that respond differently based on the genetic background of an infecting nematode. Specifically, we measured host response via gene expression profiles, and asked: can we identify variation in expression across genetically identical hosts when they are exposed to genetically variable parasites? In each host species we examined, the answer to this was yes. This indicates that while the infection process on a gross level is consistent, on a molecular level this process appears to depend on the genetic background of the parasite. The ability to connect phenotypic variation to genetic variation forms the basis for exploring how phenotype is genetically controlled. In a previous experiment (Guo et al. 2017), we were able to make specific connections between nematode genetic variants and plant (and RKN) gene expression patterns during the infection process. Extending that concept here, we were able to determine which of these patterns appeared to be conserved across widely divergent host species. This not only strengthens the finding that the infection process is highly genetically driven, but also helps direct attention towards key plant genes as being targeted by the parasite.

Interestingly, while orthologous genes across species tended to agree in terms of differential expression patterns, when genes were differentially expressed, the direction of the expression change was surprisingly inconsistent. There are several reasons that could lead to this type of outcome. One is the presence of false positive signals (yielding a mixture of true and false positives within an orthologous group). Another reason for reversed signals within an orthologous group is that, while the DNA sequence of these genes may be conserved across species, the actions of those genes may be different in the different species. A third is that the orthologous groups may not truly contain orthologous genes, or may contain multiple subgroups of orthologous genes.

As infected tissue samples (galls) were assayed for each of these experiments, we were also able to identify parasite gene expression patterns that differ according to the genetic background of the nematode. Gene expression of the nematodes was highly similar from one plant host to the other, also reinforcing the genetic nature of the infection process. Of the 93 genes that were differentially expressed between VW9 nematodes and LM nematodes for all three plant hosts, 92 were consistent in the direction of expression. Based on these results, the process by which each strain of nematode infects each of the three host plants is highly consistent, and not identical between *M. hapla* strains.

Finally, we were able to identify conserved signals between plant host species when nematode-infected root tissue was compared with uninfected tissue, in spite of the confounding factor resulting from infected and uninfected samples having been collected and assayed in different labs. Here our rationale was that the confounding factors creating false positive effects would not be expected to be conserved between tomato and soybean data. As the effects we identified were consistent between species, our results appeared to be enriched for biologically relevant differences between infected and uninfected tissue.

In all, these results demonstrate that genetic variation within *M. hapla* is a key factor in determining which plant genes are recruited for successful establishment of infection and development of feeding sites. This observation has substantial implications for the use of this system to further our understanding of these processes on a molecular level.

## ACKNOWLEDGEMENTS

Thanks to Dr. Sylwia Fudali for contributing infected Medicago tissue, and to Stella Chang for generating the infected soybean and tomato tissue. Thanks also to Dr. David Bird and Dr. Valerie Williamson for feedback on an earlier version of the manuscript. This work was funded in part by National Science Foundation grants #1025840 and #2210293.

